# Attention to a point in time causes a suppressive wake for subsequent time points

**DOI:** 10.1101/2025.04.28.651029

**Authors:** Shira Tkacz-Domb, Yaffa Yeshurun, Tony Lindeberg, John K. Tsotsos

## Abstract

The Selective Tuning model proposed that attention to a visual stimulus suppresses interfering portions of the processing hierarchy. This has been shown in spatial and featural domains and the proposal extends to the temporal dimension. We investigated whether attending to a point in time leads to processing suppression of nearby time points. We presented a sequence of letters with an embedded target, and observers indicated the target’s orientation. In the neutral condition, the target appeared randomly in one of the frames. In the informative condition, the target appeared in the same frame on most of the trials (‘expected’ trials), and the expected frame varied between blocks (Experiments 1-3) or groups (Experiment 4). On the remaining trials, it appeared before or after the most-probable frame (‘unexpected’ trials). Four different experiments were conducted to adjust the method and in the final one we found higher accuracy in expected trials than in neutral trials, indicating the allocation of temporal attention to the expected frame. When the target appeared after the expected frame, accuracy was lower than in the same neutral condition frame, suggesting an attentional suppressive wake in time (because the effect of attention cannot be observed until after stimulus onset).

## Introduction

The Selective Tuning (ST) model considers attentional ability to be comprised of a set of mechanisms whose goal is to dynamically tune perceptual processes so that they achieve their best performance for the tasks and stimuli of the moment. The three main types of attentional mechanisms are selection, restriction, and suppression (Tsotsos 2011). Selection refers to the process of selecting one element of the set of possibilities over the others, and this includes selecting a region of interest (spatial attention), selecting a feature of interest (feature-based attention), and selecting the time of interest (temporal attention). It has already been shown that both spatial and feature-based attention can manipulate visual processing to improve the perception of a selected target by suppressing interference. This capability had first been described theoretically in Tsotsos (1988, 1990) and evidence began to support this model with Caputo & Guerra (1998). Summaries appear in Carrasco (2011) and Tsotsos (2011), and much has appeared since then (e.g., Bartsch et al. 2017, Yoo et al. 2018).

According to the processing timeline of the ST model (Tsotsos et al. 2008), there are two points when such manipulation might occur. The first, occurring before a stimulus is presented, is priming due to a cue about the task to be performed, so that the task stimulus is presented to a primed architecture. The second begins at the time when a task stimulus is selected for attentive processing, thus occurring after its presentation. In the case of spatial priming, the processing network is prepared to expect a target stimulus at a particular location or region and thus the remainder of the visual field is suppressed (to some degree). For feature priming, the portions of the network selective for target irrelevant feature characteristics are suppressed. The nature of signal interference depends on stimulus context along the relevant dimensions (spatial or feature dimensions) and is determined directly by the specific architecture of the neural network (neural connectivity). In both cases, the instruction to prime is endogenous (arises due to instruction or some previously presented stimulus understood to represent a cue for a future stimulus). The application of the cue thus takes time consistent with a single top-down pass of the visual hierarchy once the instruction is understood. In both cases, the goal is to adapt the visual processing network to be ready for a particular target stimulus (consistent with the Posner cueing paradigm; Posner, 1980).

In the case of attentive selection, the goal is to improve the processing of the attended stimulus by suppressing any interfering signals. Spatially, a target stimulus is made more conspicuous by suppressing its local spatial context. In other words, any signals within the receptive fields that process this stimulus present interfering context which degrades the interpretation of the attended stimulus and their reduction or removal improves the interpretation. This is done after selection and occurs top-down throughout each layer of the visual processing network in order to suppress interfering signals from the network (both signals due to feedforward input from other stimuli and signals due to lateral inhibition within any processing layer; see Tsotsos 2011). In the feature case, interfering nearby features, that is nearby features within that feature dimension, are suppressed (Bartsch et al. 2017, Yoo et al. 2021). One might think that all targets have a spatial presence as well as other features and that these two mechanisms might interact, and this seems to be the case (Yoo et al. 2018).

We consider time to be yet a third stimulus dimension. In both spatial and feature cases, suppression reduces the effect of nearby stimuli on a target stimulus. What Cutzu and Tsotsos (2003) found is that attention to the spatial location of a stimulus modulated visual processing via suppression of spatially near stimuli presented subsequently, that is, within the processing receptive fields of the target throughout the network. Since this observation has been repeated many times for the spatial and featural domains, we presumed that this might apply also to the temporal domain, thus the goal of our study. Specifically, an observer could modulate processing structures across time in anticipation of a target so that they are optimally set for its detection and interpretation.

A straightforward generalization of past results might say that the choices would be to enhance the expected target and/or its processing, to suppress some or all of the other stimuli or their processing, or some combination of these. However, the temporal domain has distinct differences. Although spatially near stimuli within a single image (and this applies for stimuli nearby in featural or object dimensions) remain accessible during all stages of the processing of that image, temporally ordered stimuli necessarily overwrite the preceding ones as they flow through the visual processing system. One could expect an onset at a particular point in time and hope to suppress any competing onsets. However, “nearby” in time means earlier as well as later stimuli. Once a target is detected the preceding stimuli are not so directly accessible, no actions are possible in any previous time step because one cannot affect the past. Could neurons be primed to enhance their sensitivity at a future time? Without a clear mechanistic way to modulate neural circuits in the future it is difficult to see how this might be possible. Would there be a ‘clock’ to provide the timing of when to modulate? There is little evidence for such a mechanism. This leaves the possibility for attentive modulation in the future, i.e., after the stimulus appears.

Imagine a steady stream of stimuli passing through the visual system, much like a constant wave. This, of course, is the reality of real-world vision; the flow of visual stimuli into the eyes is continuous and the real world is dynamic and ever-changing. What could be an appropriate probe into this stimulus stream? It might be useful to think of a fruit processing factory where fruits are continuously dumped onto a conveyor belt and pass by inspectors at a constant rate. Each stimulus (or observation of a single fruit) has a very short duration and its existence within the observer’s system is limited to the time of passage through the visual system of that fruit ‘wave’ because it is immediately replaced in the wave by other fruit (recall how it was shown that visual category decisions can be made in about 150ms, Thorpe et al., 1996; the stimulus wave in our fruit example has a similar lifetime within the visual system).

Let’s say there is an expectation for a stimulus at time *t*. Any modulation due to the expectation could occur before, during or after time *t*, and would have some duration *k.* Suppose attention suppresses stimuli in advance of the expected time, say from *t-k* to *t*. This would make the onset of stimuli at *t* more prominent. But without any spatial expectation the suppression would be for the whole image and then all stimuli would be seen as onset at time *t* giving the target no advantage. Even if such an additional spatial expectation is given, the observer would see no temporal modulation within the expected spatial area, and again there is no advantage due to the temporal prior. On the other hand, suppose suppression occurs for stimuli that follow the expected time point *t*, from *t* to *t+k*. To be clear, any stimuli that precede the target time would still exist but at later stages of the processing network while those appearing after the target time would be found in earlier stages of processing.

Thus, suppression of the following stimuli means that any interference during the interpretation of the target is subdued - the subsequent stimuli are prevented from overwriting the stimulus of interest for a short time - leading to a clearer decision about the stimulus seen at the expected time point. In other words, if the expected piece of fruit is followed by a blank conveyor belt, even for a short time, one can complete the processing of that fruit stimulus more effectively. We term this the *suppressive wake* that accompanies attention in time, and our experiments seek to reveal evidence for it.

Lindeberg has presented a model of early visual processing that includes the temporal component with a strong resemblance to what we have described in actual neurons (Lindeberg 2021). The tuning profile in time consists of a positive region followed by a negative one. The overall model is very favorably compared to actual retinal, LGN, and V1 neurons in mammals. The figure below (from Figure 10, Lindeberg 2021) shows the receptive field profiles in the x-dimension (Figure 1a - a positive and a negative Gaussian, as seen in an edge-selective neuron) and in the t-dimension (Figure 1b - a positive Gaussian followed by a negative one) to illustrate. On the surface this seems to mean that this neuron has a preference for an edge-like stimulus of short duration in time. However, this might have a secondary meaning as well. The suppression of future signals (the negative lobe) makes the current one (the positive lobe) more conspicuous as the signal traverses through higher layers of visual processing. This seems an important mechanistic aspect for the attentive temporal suppression we are hypothesizing.

**Figure 1.**
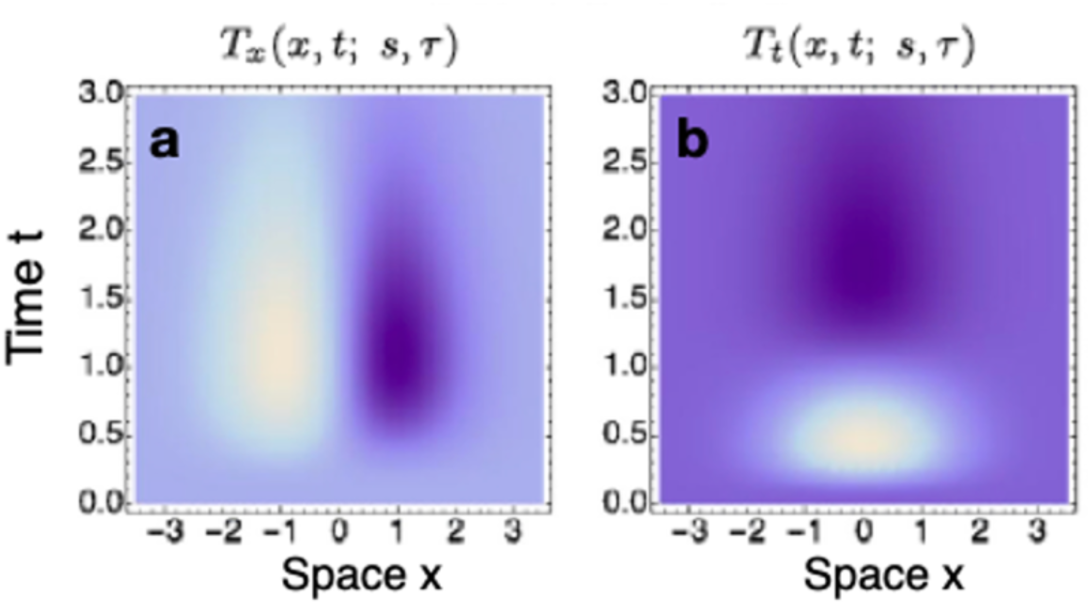
Illustration of space-time separable receptive fields *T_x_^m^ _t_^n^ (x,t; s,τ)=* *∂_x_ ^m^ _t_^n^(g(x; s)h(t; *τ*))* of order one. *s*=1, *τ*=1. Figure reprinted with permission from Lindeberg 2021.

In the neurons Lindeberg (2021) describes, these characteristics are built in; they are not attentive but part of the inherent tuning curve of the neurons. Many have shown attentive manipulations of neural properties throughout the visual processing network from LGN to all higher layers (Mehta et al., 2000; O’Connor et al., 2002). The manipulations all affect the basic neural tuning properties (such as those in Figure 1a) for spatial (e.g., Fischer & Whitney, 2009; Roberts et al., 2007; Womelsdorf et al., 2006) and feature dimensions (e.g., David et al., 2008; Haenny, & Schiller, 1988; Martinez-Trujillo, & Treue, 2004). Our question then asks could the same kinds of attentional manipulations be seen in the temporal dimension, for the tuning shown in Figure 1b? The prediction that guides the current research then, following from the past demonstration of suppression in the spatial and feature dimensions and strongly supported predictions of the Selective Tuning model, is that an attentive suppression will be seen in time and our experiments test this for the priming scenario described earlier.

Typically, orienting endogenous temporal attention has been manipulated in experimental tasks using symbolic cues (e.g., the word “early” or “late”) that indicate the time interval after which the target is most likely to appear. The cue can be either valid (“early” for a short interval) or invalid (“late” for a short interval), and the typical result is better performance in valid than invalid trials (e.g., Griffin et al., 2001; Yeshurun & Tkacz-Domb, 2021). Another prevalent way to manipulate endogenous temporal attention is by using hazard rates. Changes in the probability of the time point in which the target appears after a warning signal (“ready”) modulate performance (e.g., Cravo et al., 2011; Ghose & Maunsell, 2002). For example, when the target is equally likely to appear at any of a number of possible times (uniform distribution), performance increases as the waiting period between the warning signal and target appearance is longer. It is assumed that as a longer waiting period elapsed, the temporal uncertainty reduces, and the probability that the target will appear rises, given that it has not yet appeared (e.g., Baumeister, & Joubert, 1969; Niemi & Näätänen, 1981).

In this study, we manipulated the allocation of temporal attention by modifying the probability of target appearance in the different conditions. Unlike previous studies in which the target was the only stimulus that appeared after the warning signal (e.g., Cravo et al., 2011), our target appeared within a sequence of stimuli. Specifically, we designed a task in which a target letter was embedded in a long sequence of other letters, and observers were asked to indicate its orientation. In the neutral condition, the target appeared randomly and with equal probability in one of the frames in the sequence, so that target timing was not predictable. In the informative condition, the target appeared in the same frame within the sequence on most of the trials (‘expected’ trials). On the rest of the trials, the target appeared before or after the most-probable frame (‘unexpected’ trials). In three experiments, the most-probable frame varied between blocks, and the observers were told which is the most-probable frame at the beginning of the block. This information allowed participants to estimate when the target most often appeared and volitionally allocate temporal attention to that moment. In the fourth experiment, five different groups of participants conducted different conditions. Four groups participated in the informative condition, and the most-probable frame was different in each group. The fifth group participated in the neutral condition. If orienting voluntary temporal attention to a particular moment leads to enhancement at the attended time point as well as suppression at succeeding time points, performance should be higher when the target appears in the expected frame than when it appears in the same frame in the neutral condition, and it should be lower when the target appears in an unexpected frame after the anticipated time than in the same frame in the neutral condition. Thus, our hypothesis predicts an orderly movement of the suppression that tracks the expected time point.

## Experiment 1

## Methods

### Observers

Sixty students from York University participated in this experiment, 6 of whom were excluded from the analysis due to poor performance (3 participants had accuracy below 50%, 2 participants had accuracy at guessing level in 3 blocks, and 1 participant did not follow the instructions and sat too close to the screen). All observers had normal or corrected to normal vision and were naïve to the purpose of the study. Power analysis using G*Power 3.1 (Faul et al., 2007) and the effect size reported by a study that examined the attentional suppressive surround in both the spatial dimension and feature dimension (Yoo et al., 2018) suggested we would need 20 observers for examination of suppression effects at a power >95 % with a Type 1 error of α < 0.05. Because the experiment was conducted online, and the observers’ environment tended to change more than in the lab (e.g., light, noise, etc.), we recruited a larger group of participants to reduce noise in the data. This study adhered to the Declaration of Helsinki, and was approved by the ethics committee of York University (certificate number 2020-029). Observers provided informed consent and received course credit or payment for participation.

### Apparatus and stimuli

The experiment was created in PsychoPy (version 2020.2.4; Peirce et al., 2019) and hosted on the online platform Pavlovia. Observers’ monitor refresh rate was 60 Hz. The fixation mark was a central black plus (0.2° x 0.2°). A sequence of 10 rotated letters (A C E H L M P S U T; 1° x 1°; RGB [88 88 88]) appeared at the center of the screen (Figure 2). One of these letters was the target T, which could be upright, inverted, turned left, or turned right. The observers had to indicate its orientation by pressing the corresponding key on the keyboard. Each letter was displayed on a square (1.2° x 1.2°) of static random-dot noise. The background was a uniform gray (RGB [128 128 128]).

**Figure 2.**
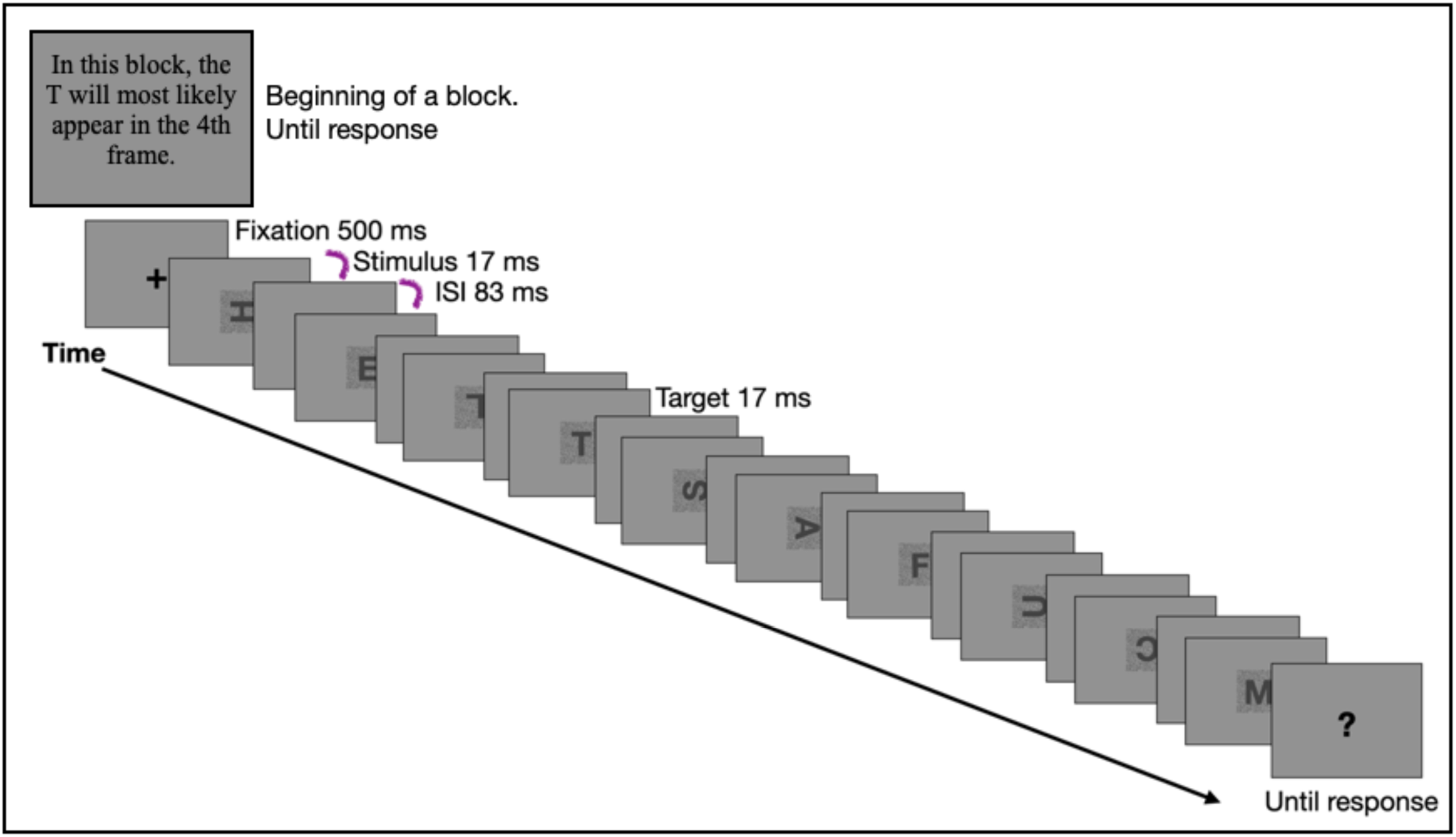
A schematic example of a trial in the informative block in Experiment 1. In this example, the target (the letter T) is expected to appear in the 4th frame, and it can appear unexpectedly in the 2nd, 3rd, 5th, or 6th frames. Each letter in the stream appeared for 17 ms, and between letters, there was an inter-stimulus-interval (ISI) in which a blank display appeared for 83 ms.

### Procedure

To control for stimulus size, before beginning the experiment, we estimated the screen resolution of the observers’ monitor and their distance from it by using the Virtual Chinrest method described by Li et al. (2020). Using this information, we presented the stimuli to all observers at the same size in visual angles. After completing this task, the experiment began. At the beginning of each block, participants were told whether the upcoming block was informative or neutral. On informative blocks, in 75% of the trials, the target appeared in the same frame within the sequence (‘expected’ condition). In 25% of the trials, the target appeared one or two frames before or after the most-probable frame (‘unexpected’ condition). The most-probable frame varied between blocks (4th, 5th, 6th, or 7th frame). To help participants learn the expected time point, they were told which is the most-probable frame at the beginning of each block. In the neutral block, the target appeared randomly in one of the frames (frames 2-9). Each trial started with a central fixation mark. After 500 ms, the sequence of letters appeared. The letters were separated by a stimulus-onset asynchrony (SOA) of 100 ms (stimulus presentation duration was 17 ms and inter-stimulus-interval was 83 ms). Once the sequence was over, observers had to indicate the orientation of the target T. Then, visual feedback appeared at the screen’s center (correct: green “+”; incorrect: red “-“). The experiment consisted of 2 sessions. Each session included two different informative blocks, each with 320 experimental trials. The informative blocks that were chosen for the different sessions were counterbalanced across observers. The neutral block included 160 experimental trials, and it always appeared in the first session. Blocks’ order was random within each session. Fifty practice trials preceded each block, and observers had a break every 40 experimental trials. The second session was available after 2 hours from the end of the first session and up to 72 hours.

## Results

To analyze the data of the 54 participants, we first conducted a 1-way (expectancy) repeated-measures analysis of variance (ANOVA) on the accuracy data. This analysis revealed a significant main effect of expectancy (F(2,106)=18.9, *p*<0.0001, ƞ_p_^2^=0.26), accuracy was highest for expected trials (80%), intermediate for unexpected trials (78%), and lowest for neutral trials (75%). We note that in this analysis, the unexpected condition includes unexpected target positions for which attentional suppression was not anticipated because these positions preceded the expected position. Because there were different target positions in the different informative blocks, performing an omnibus ANOVA that includes all the different informative and neutral blocks was impossible. Thus, we conducted a separate 2-way repeated-measures ANOVA with the factors informativeness (informative vs. neutral) and target position for each of the informative blocks (Figures 3 and 4). A significant main effect of target position emerged in all of the four analyses (4th frame expected: F(4,212)=12, *p*<0.0001, ƞ_p_^2^=0.2; 5th frame expected: F(4,212)=17.3, *p*<0.0001, ƞ_p_^2^=0.24; 6th frame expected: F(4,212)=13.6, *p*<0.0001, ƞ_p_^2^=0.2; 7th frame expected: F(4,212)=19.5, *p*<0.0001, ƞ_p_^2^=0.27). A marginal interaction was found in informative blocks in which the expected frame was the 4th and 5th (F(4,212)=2.03, p=0.09, ƞ_p_^2^= 0.04; F(4,212)=2.1, p=0.08, ƞ_p_^2^= 0.04, respectively). Least significant difference (LSD) post hoc tests indicated that accuracy was significantly higher when the target appeared in the expected frame rather than the same frame in the neutral condition (Figure 4). This suggests that attention was allocated to the expected frame. Although accuracy decreased after the expected frame, it was not significantly lower than in the neutral condition. Therefore, there was no evidence of suppression in time.

**Figure 3.**
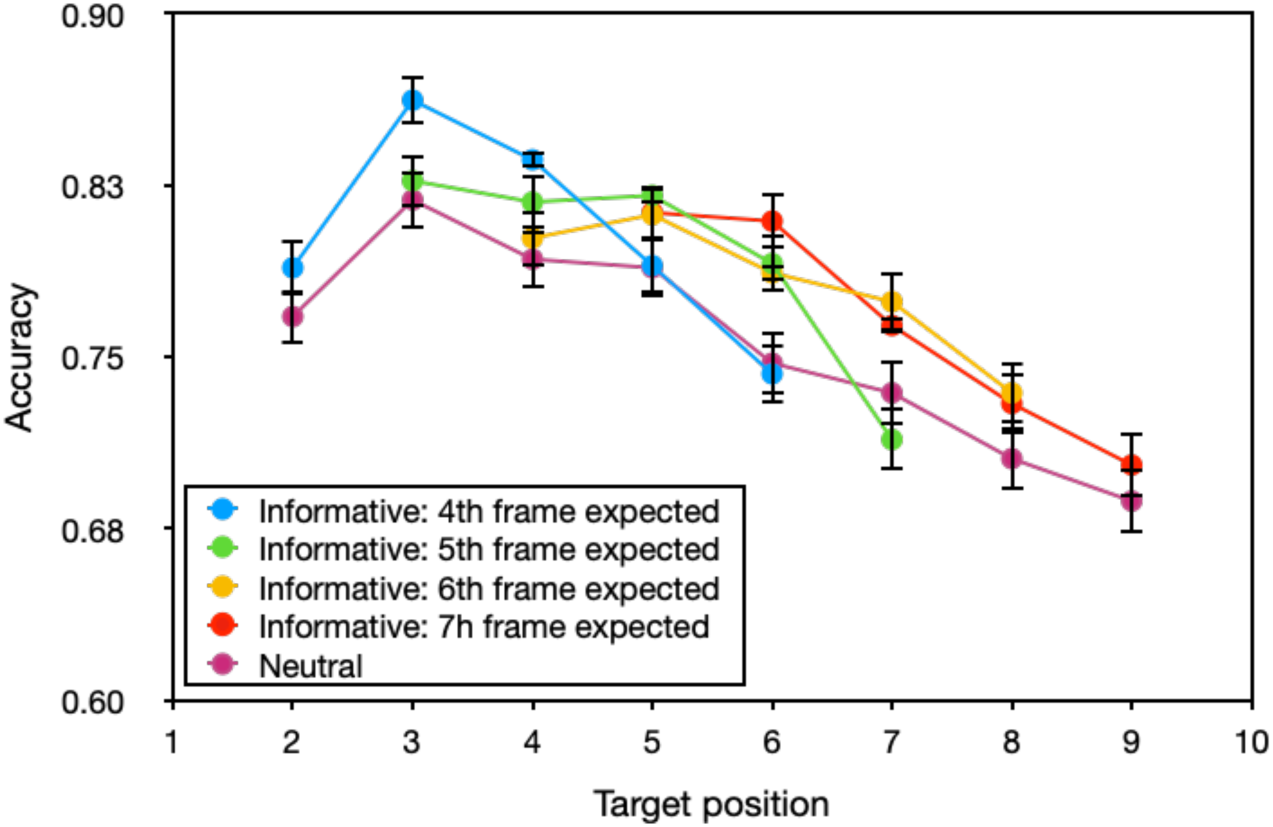
Accuracy in Experiment 1 as a function of informativeness and target position in the sequence of stimuli. Error bars correspond to one standard error of the means (SEM) across observers.

**Figure 4.**
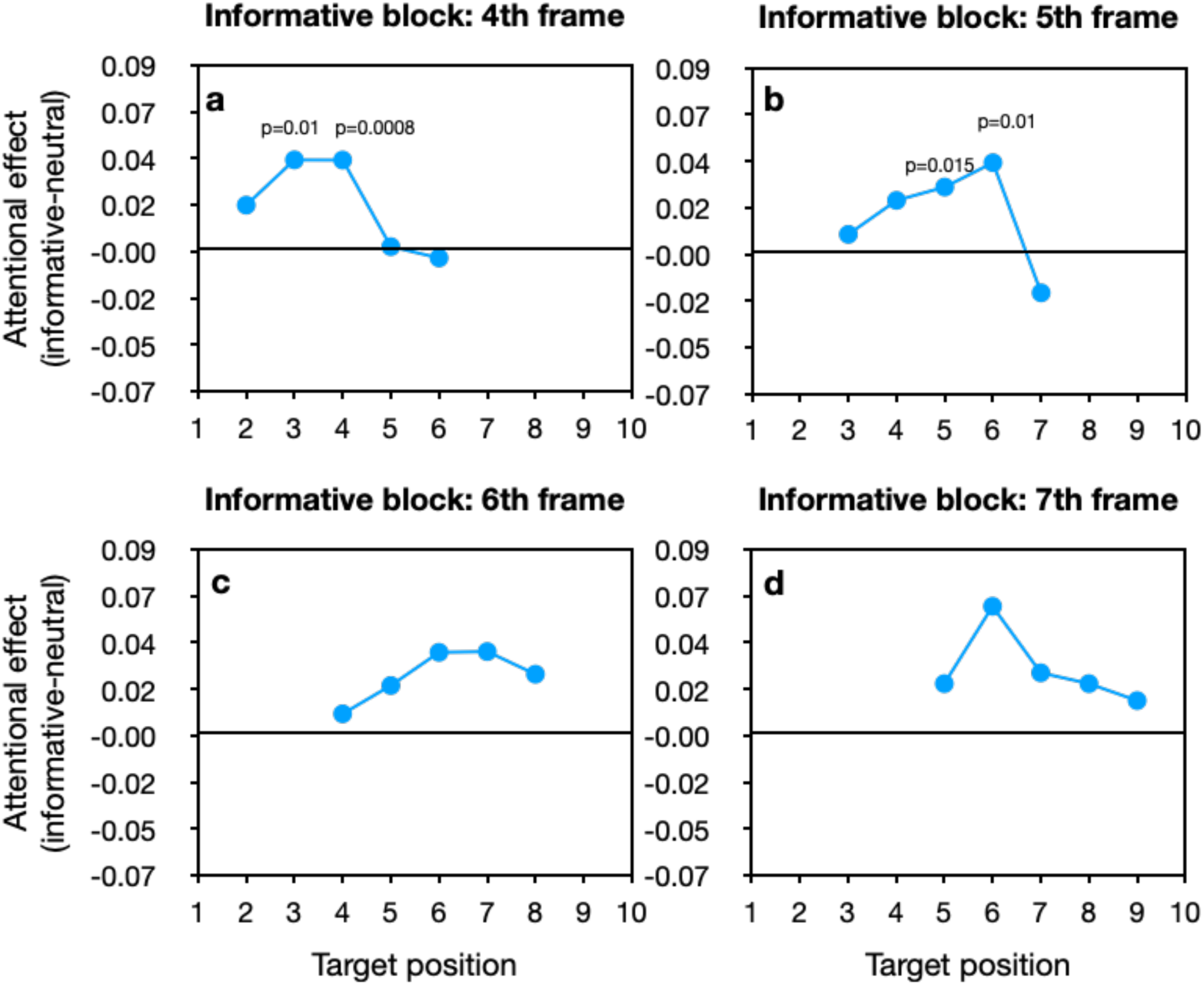
The attentional effect in Experiment 1 as a function of target position within the stream. (a) informative condition in which the 4th stimulus position was the expected frame; (b) informative condition in which the 5th stimulus position was the expected frame; (c) informative condition in which the 6th stimulus position was the expected frame; (d) informative condition in which the 7th stimulus position was the expected frame. The p-value for each LSD post-hoc test of the difference between the neutral block and an informative block for a given target position is presented above that target position.

## Experiment 2

In Experiment 1, we found a decrease in performance after the expected time, but it was not significantly lower than the neutral condition. Experiment 2 tested the possibility that the suppression requires more time to develop than the time points tested in Experiment 1 and would therefore be revealed after a longer interval from the expected time point. To that end, Experiment 2 was similar to Experiment 1, but the target could also appear three frames after the anticipated frame.

## Methods

### Observers Stimuli and Procedure

Sixty-two observers participated, of which 19 were excluded from the analysis due to poor performance (13 of them had an accuracy of less than 50% over the entire experiment, 4 participants had an accuracy below 50% in one of the sessions, and 2 participants had an accuracy below 50% in 6 or more blocks towards the end of the experiment). Five of them also participated in Experiment 1. The experiment was similar to Experiment 1, except for the following: First, the target could appear up to three frames after the expected frame. Second, the sequence included 13 letters (A C E H L M P S U F R Y T). Third, in the neutral condition, the target could appear in one of the frames from the 2nd to the 12th. Finally, each informative block included 400 experimental trials and was preceded by 40 practice trials. The neutral block included 220 experimental trials and was preceded by 44 practice trials. The second session was available after 2 hours and up to 96 hours from the end of the first session.

## Results

Similar to Experiment 1, a 1-way repeated-measures ANOVA showed the highest accuracy for expected trials (78%), intermediate for unexpected trials (76%), and lowest for neutral trials (71%), F(2,84)=43.5, *p*<0.0001, ƞ_p_^2^=0.5. In addition, a separate 2-way (informativeness x target position) repeated-measures ANOVA for each informative block, revealed in all analyses, a significant effect of target position (4th frame expected: F(5,210)=5.1, *p=*0.0002, ƞ_p_^2^=0.1; 5th frame expected: F(5,210)=6.5, *p*<0.0001, ƞ_p_^2^=0.13; 6th frame expected: F(5,210)=9.4, *p*<0.0001, ƞ_p_^2^=0.18; 7th frame expected: F(5,210)=13.7, *p*<0.0001, ƞ_p_^2^=0.24). The interaction of informativeness and target position was significant when the expected frame was the 4th and 5th (F(5,210)=2.4, *p*=0.04, ƞ_p_^2^=0.05; F(5,210)=2.2, *p*=0.05, ƞ_p_^2^=0.05, respectively), and marginally significant when it was the 6th frame (F(5,210)=1.9, *p=*0.09, ƞ_p_^2^=0.04), (Figures 5 and 6). Further LSD post-hoc tests showed significantly higher accuracy around the expected frame than the same frame in the neutral condition, suggesting that attention was allocated to the expected frame. This was the case with all informative blocks, apart from the 7th frame informative block. Accuracy decreased after the expected frame, but not below the neutral condition. One possible reason for the lack of evidence for suppression could be the unexpected accuracy decline with target position in the neutral condition (i.e., a decline observed even though in this condition all target positions are equally likely). We wondered whether this decline is related to carryover expectations when the neutral condition followed an informative condition. To test this hypothesis, we conducted a 2-way mixed-design ANOVA on the accuracy in the neutral condition, with target position as a within-subject variable and block order as a between-subject variable. This analysis revealed a significant interaction (F(20,400)=3.2, *p<*0.0001, ƞ_p_^2^=0.14; Figure 7b). The effect of target position was not significant when the neutral condition was the 1st block in the session (F(10,150)=0.7, *p*=0.7, ƞ_p_^2^=0.05), but was significant when it was the 2nd or 3rd block, (F(10,140)=4.7, *p*<0.0001, ƞ_p_^2^=0.25; F(10,110)=6.6, *p*<0.0001, ƞ_p_^2^=0.4, respectively). This interaction was not significant in Experiment 1 (F(14,357)=1.1, *p*=0.4, ƞ_p_^2^=0.04; Figure 7a). Still, for the sake of comparison, we also performed separate analyses of the target position for each order of the neutral block in Experiment 1. Once again, the effect of target position was not significant when the neutral condition was the 1st block (F(7,98)=1.3, *p*=0.3, ƞ_p_^2^=0.08), only when it was the 2nd or 3rd block in the session (F(7,175)=4.9, *p*<0.0001, ƞ_p_^2^=0.17; F(7,84)=4.7, *p=*0.0002, ƞ_p_^2^=0.28). Because the neutral block was only displayed in the first session, when it was the first block, the observers had not yet been exposed to any informative block and had no reason to adopt any expectations regarding the target position. In contrast, when the neutral block followed informative blocks, in which the target did not appear after the 7th position, accuracy decreased in target positions larger than 7, possibly due to learned probabilities that were carried over from previous informative blocks. Hence, the lack of suppression may be due to carryover effects from the informative blocks to the neutral block. In addition, in both experiments, the accuracy was higher in the second session than the first (Experiment 1: F(1,53)=16.5, p=0.0002, ƞ_p_^2^=0.24; Experiment 2: (F(1,42)=9.5, p=0.004, ƞ_p_^2^=0.19). We compared the informative blocks, which appeared in both sessions, with the neutral block, which appeared only in the first session. Therefore, if the higher accuracy in the second session was due to improved task performance, then the lack of finding suppression may also be because of lower accuracy in the neutral due to less training in the task.

**Figure 5.**
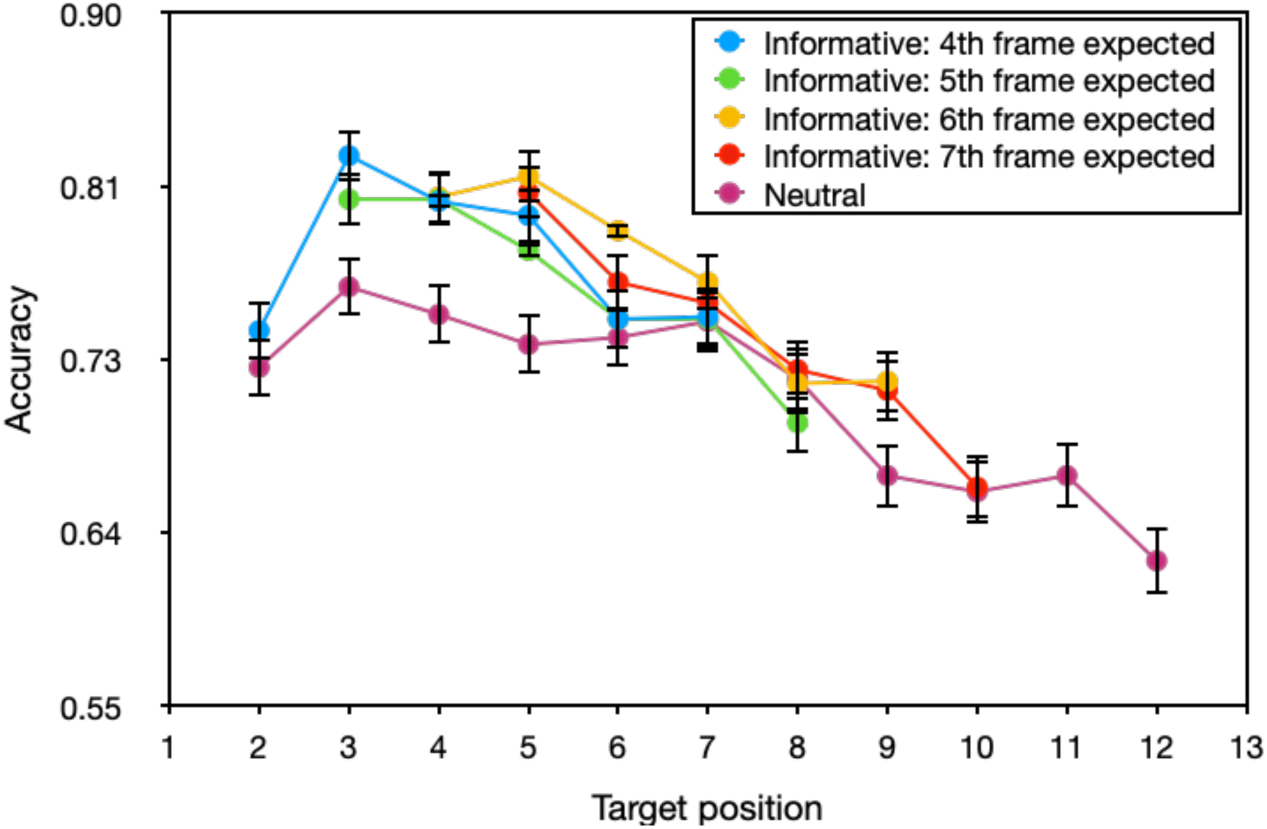
Accuracy in Experiment 2 as a function of informativeness and target position in the sequence of stimuli. Error bars correspond to one SEM across observers.

**Figure 6.**
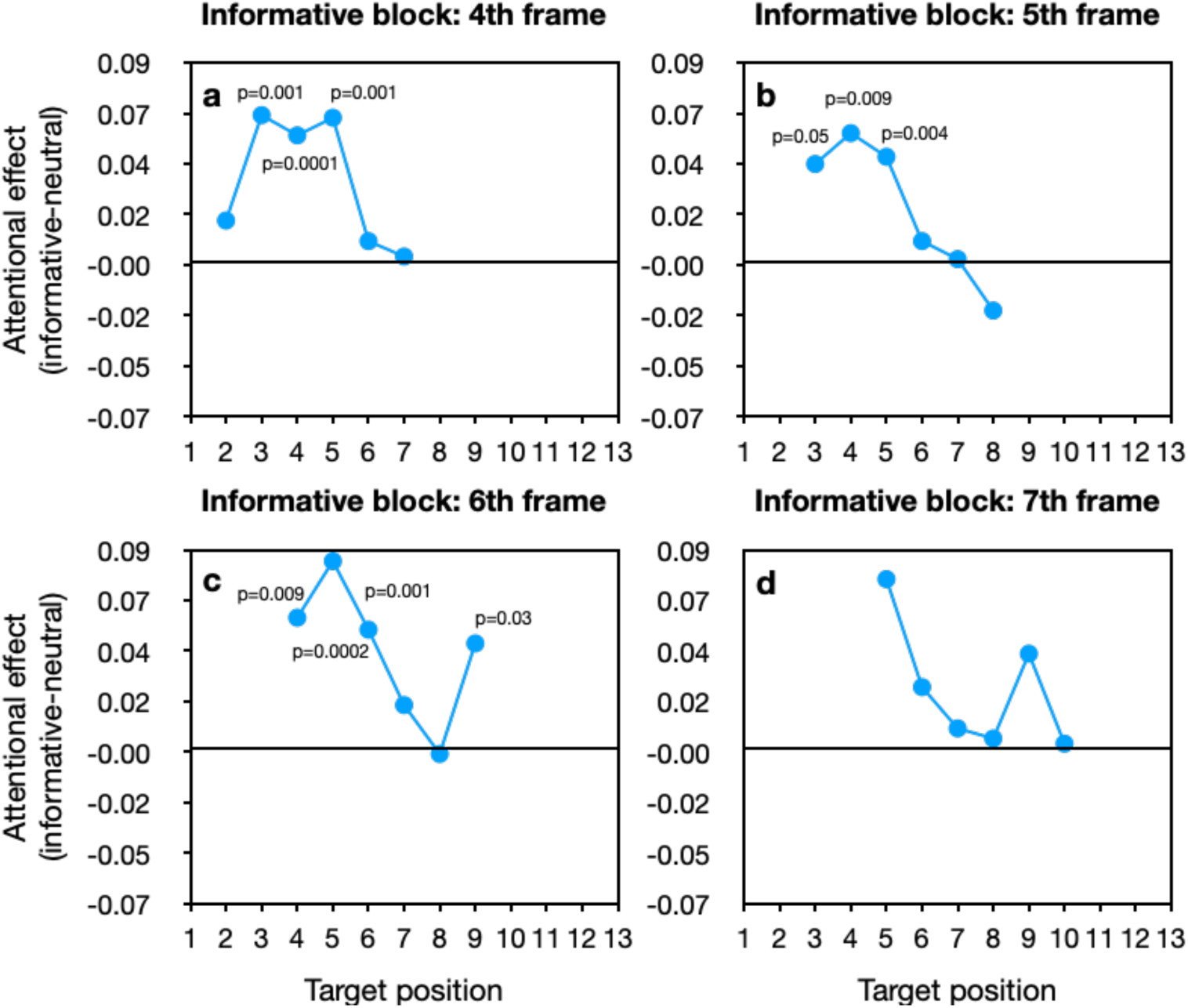
The attentional effect in Experiment 2 as a function of target position within the stream. (a) informative condition in which the 4th stimulus position was the expected frame; (b) informative condition in which the 5th stimulus position was the expected frame; (c) informative condition in which the 6th stimulus position was the expected frame; (d) informative condition in which the 7th stimulus position was the expected frame. The p-value for each LSD post-hoc test of the difference between the neutral block and an informative block for a given target position is presented above that target position.

**Figure 7.**
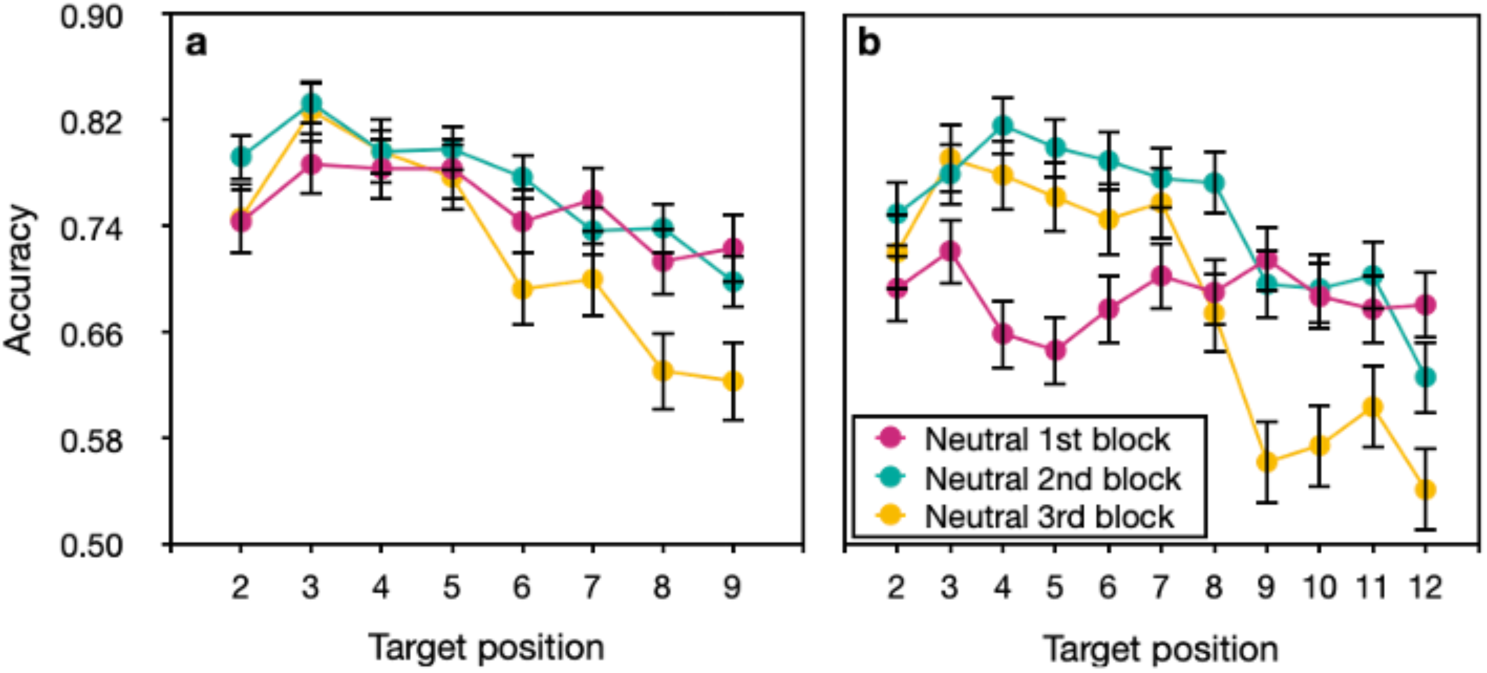
Accuracy as a function of neutral block order and target position within the stream in (**a**) Experiment 1 and (**b**) Experiment 2. Error bars correspond to one SEM across observers.

## Experiment 3

In Experiments 1 and 2, we did not find suppression in time, but the lack of suppression may result from carryover effects from the informative blocks to the neutral block. To overcome these carryover effects, in Experiment 3, the neutral block was always the first block in the session, and it appeared in both sessions.

## Methods

### Observers Stimuli and Procedure

Seventy-one observers participated, 15 of whom were excluded from analysis due to poor performance (8 participants had accuracy below 50% in one of the sessions, 5 participants had accuracy below 50% in more than 6 blocks towards the end of the experiment, and 2 participants sat too close to the screen). Two participants were recruited through Prolific and received payment. The rest were students from York University who received course credit. The experiment was similar to Experiment 2 except for the following: The sequence consisted of 11 letters (A C E H L M P S U F T). In the neutral condition, the target could appear in one of the frames from the 2nd to the 10th. The neutral block was the first block in both sessions. Each informative block included 400 experimental trials and was preceded by 40 practice trials. The neutral block included 360 experimental trials and was preceded by 45 practice trials. The second session was available after 4 hours from the end of the first session and up to 96 hours, except for one participant from Prolific who performed the second session after a month.

## Results

Similar to the previous Experiments, accuracy was higher in the expected condition (80%) compared to the neutral (76%) and unexpected (77%) conditions (F(2,110)=30.7, p<0.0001, ƞ_p_^2^=0.36). In addition, the accuracy increased in the second session (F(1,55)=38.4, p<0.0001, ƞ_p_^2^=0.4). We performed a separate 2-way (informativeness x target position) repeated measures ANOVA for each of the four informative blocks. All analyses revealed a significant effect of target position (4th frame expected: F(5,275)=13.8, p<0.0001, ƞ_p_^2^=0.2; 5th frame expected: F(5,275)=15.7, p<0.0001, ƞ_p_^2^=0.2; 6th frame expected: F(5,275)=19, p<0.0001, ƞ_p_^2^=0.26; 7th frame expected: F(5,275)=31.9, p<0.0001, ƞ_p_^2^=0.37) and interaction (4th frame expected: F(5,275)=2.7, p=0.02, ƞ_p_^2^=0.05; 5th frame expected: F(5,275)=3.8, p=0.002, ƞ_p_^2^=0.06; 6th frame expected: F(5,275)=4.8, p=0.0003, ƞ_p_^2^=0.08; 7th frame expected: F(5,275)=2.9, p=0.01, ƞ_p_^2^=0.05), (Figures 8 and 9). Additional analyses with LSD post-hoc tests revealed that when the expected frame was the 5th or the 6th, accuracy was significantly higher in the expected frame than in the same frame in the neutral block, and it decreased afterwards.

**Figure 8.**
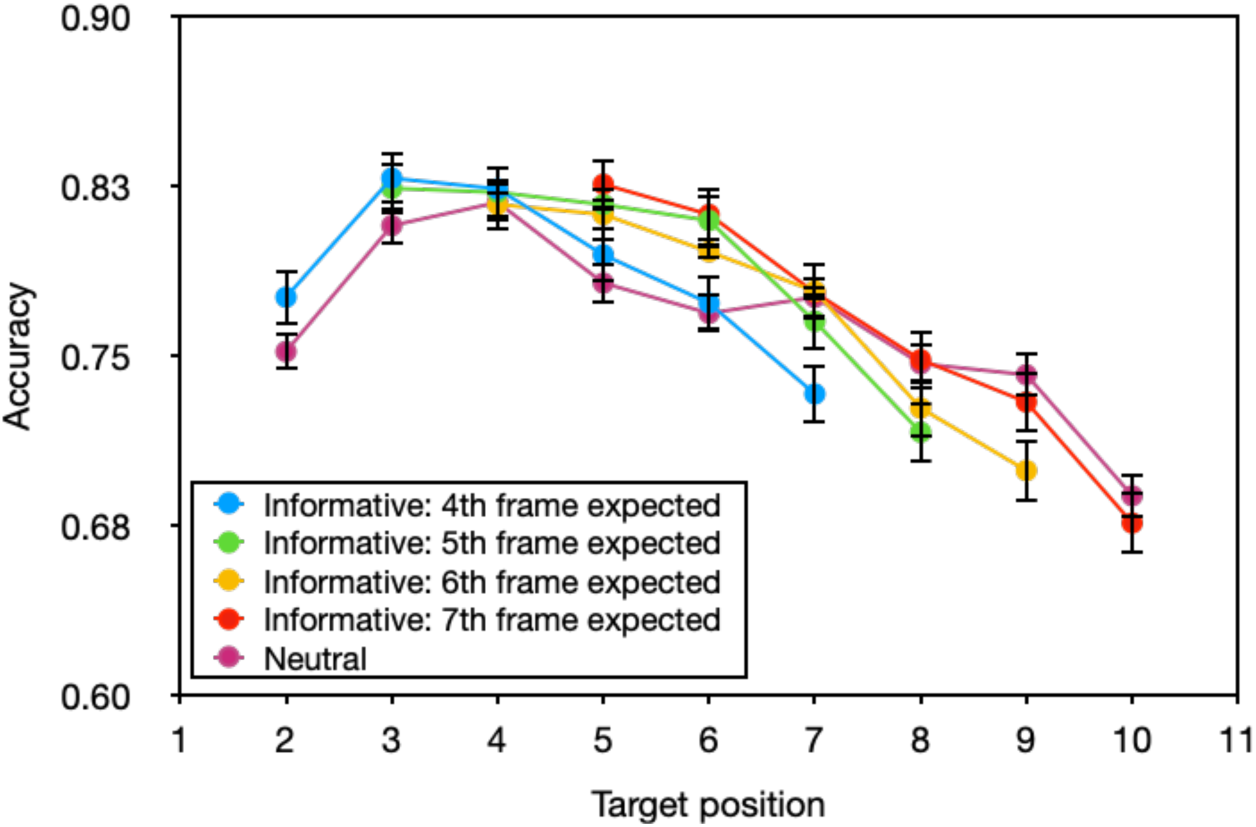
Accuracy in Experiment 3 as a function of informativeness and target position in the sequence of stimuli. Error bars correspond to one SEM across observers.

**Figure 9.**
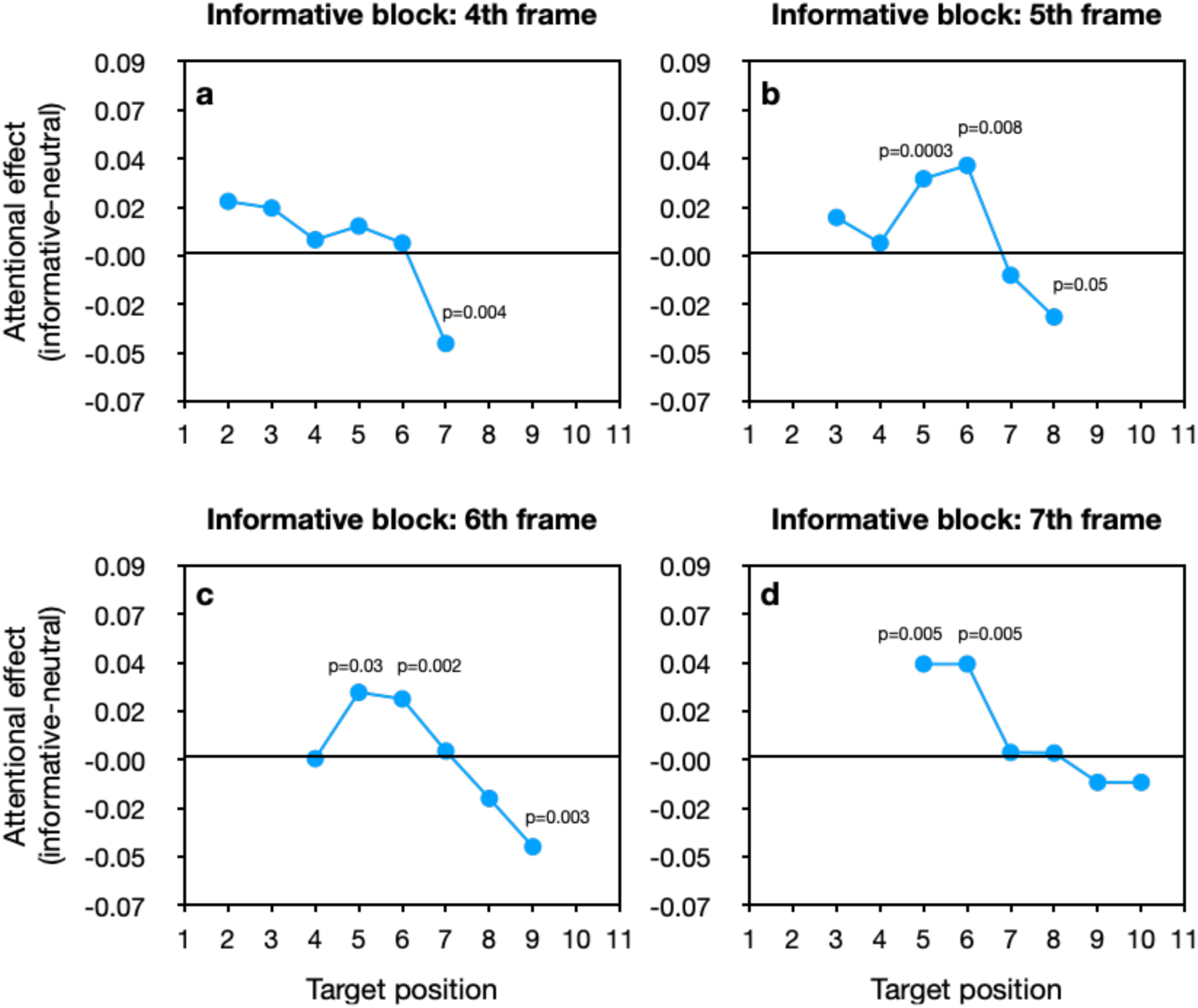
The attentional effect in Experiment 3 as a function of target position within the stream. (a) informative condition in which the 4th stimulus position was the expected frame; (b) informative condition in which the 5th stimulus position was the expected frame; (c) informative condition in which the 6th stimulus position was the expected frame; (d) informative condition in which the 7th stimulus position was the expected frame. The p-value for each LSD post-hoc test of the difference between the neutral block and an informative block for a given target position is presented above that target position.

Moreover, when the target appeared three frames after the expected frame, accuracy was lower in this unexpected frame compared to the same frame in the neutral block. These findings demonstrate that allocating attention to the expected frame could be accompanied by suppression after this frame. To examine whether performance differed in the neutral condition between the sessions, we performed 2-way (target position x session) repeated measures ANOVA on the accuracy data in the neutral condition (Figure 10). Accuracy was higher in the second session (F(1,55)=23.1, p<0.0001, ƞ_p_^2^=0.3), and it decreased as the target appeared towards the end of the sequence (F(8,440)=16.2, p<0.0001, ƞ_p_^2^=0.23). Furthermore, a significant interaction revealed different patterns of target position effect in the different sessions (F(8,440)=2.4, p=0.01, ƞ_p_^2^=0.04). This interaction suggests that block order still influenced performance, and so the current design did not completely overcome the problem of carryover effects.

**Figure 10.**
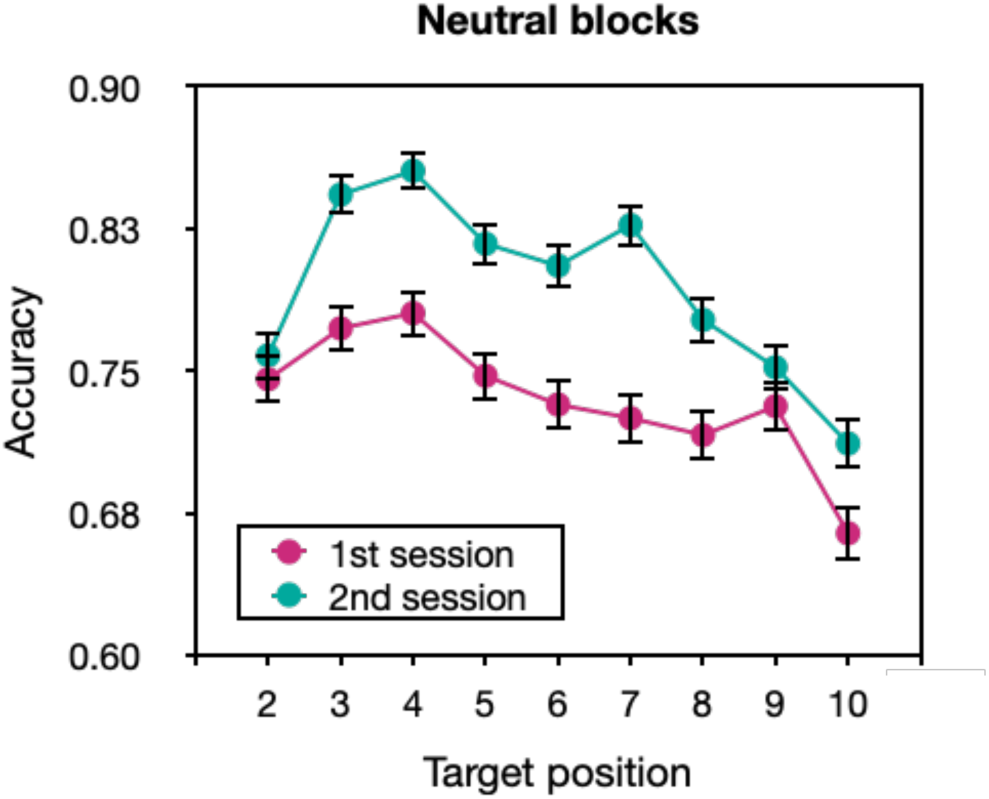
Accuracy as a function of session and target position for the neutral condition in Experiment 3. Error bars correspond to one SEM across observers.

## Experiment 4

In Experiment 3, we found that performance in the neutral condition differed in the first and second sessions, suggesting that there may still be a transfer of expectations between informative blocks from the first session to the neutral block in the second session. Therefore, Experiment 4 was designed to completely overcome the problem of carryover effects between the different conditions. Thus, each participant was exposed to only one condition in a complete between-participants design.

## Methods

### Observers Stimuli and Procedure

Two hundred and seventy one observers participated in this experiment, 21 of whom were excluded from the analysis due to accuracy below 44%. All were recruited via Prolific and received payment. The experiment was similar to Experiment 3 except that the 250 participants (after exclusion) were divided into five groups of 50 participants each. Each group performed a different condition. Four groups participated in the informative condition, with a different expected frame for each group: the 4th, 5th, 6th, or 7th frame. The additional group participated in the neutral condition.

## Results

First, we conducted a 2-sample one-sided t-test with Benjamini-Hochberg correction between the neutral group and the expected condition in the informative groups. This analysis revealed higher accuracy in the expected condition (77%) than in the neutral condition (74%) (t(248)=-1.94, p=0.03, Cohen’s d=0.3). Likewise, a one-sided paired t-test with Benjamini-Hochberg correction between the expected condition and the unexpected condition in the informative groups revealed higher accuracy in the expected than in the unexpected condition (74%), (t(199)=7.49, p<0.0001, Cohen’s d=0.5). However, a significant difference between the neutral group and the unexpected condition in the informative groups did not emerge (t(248)=-0.2, p=0.4, Cohen’s d=0.04) in a 2-sample one-sided t-test with Benjamini-Hochberg correction.

Next, we compared the results of each group that performed an informative condition to those obtained in the neutral group. We performed a 2-way mixed-design ANOVA on the accuracy for each comparison. The between-subjects factor was group and the within-subject factor was target position (Figures 11 and 12). A main effect of target position and interaction emerged in all four analyses (group of 4th frame vs. neutral group: F(5,490)=14.14, p<0.0001, ƞ_p_^2^=0.13, F(5,490)=8.7, p<0.0001, ƞ_p_^2^=0.08; group of 5th frame vs. neutral group: F(5,490)=14.4, p<0.0001, ƞ_p_^2^=0.13, F(5,490)=5.9, p<0.0001, ƞ_p_^2^=0.06; group of 6th frame vs. neutral group: F(5,490)=12.7, p<0.0001, ƞ_p_^2^=0.11, F(5,490)=4.7, p=0.0003, ƞ_p_^2^=0.05; group of 7th frame vs. neutral group: F(5,490)=13.3, p<0.0001, ƞ_p_^2^=0.12, F(5,490)=3.35, p=0.005, ƞ_p_^2^=0.03). LSD post-hoc analyses showed that in all the informative groups, the accuracy was higher around the expected frame compared to the same frame in the neutral group (for all p<0.05, except when the 7th frame was expected). Furthermore, when the target appeared after the expected frame, the accuracy decreased, even below the same frame in the neutral group (for all p<0.05). These findings suggest that allocating temporal attention to a specific time point is followed by suppressing other time points. Thus, once carryover effects were ruled out, suppression effects emerged. The results show an orderly movement of the suppression that follows the expected time, but it is a bit noisy. Specifically, when the 4th frame was expected and when the 5th frame was expected, the suppression started at the same time, in the 6th frame. This might be due to variability in the speed of processing within each subject’s visual systems versus the fixed SOA. It is also possible that the SOA we chose was not sensitive enough to the differences between positions 4 and 5, and from the participants’ point of view, these trials predicted the same time for target appearance. In addition, as seen in Figure 12, the pattern of the attentional effect, a positive effect followed by a negative one, qualitatively matches Figure 1b. In Figure 1b, in the time dimension, the receptive field is characterized by a positive Gaussian followed by a negative one. Likewise, Figure 12 also matches one side of the Mexican-hat profile that many studies have shown in both the spatial dimension and the feature dimension (e.g., Cutzu & Tsotsos, 2003; Müller et al., 2005; Tombu, & Tsotsos, 2008; Yoo et al., 2018). Interestingly, as shown in Figures 11 and 12, performance benefit arose also when the target appeared before the expected frame, indicating that the participants allocated attention before the predicted time.

**Figure 11.**
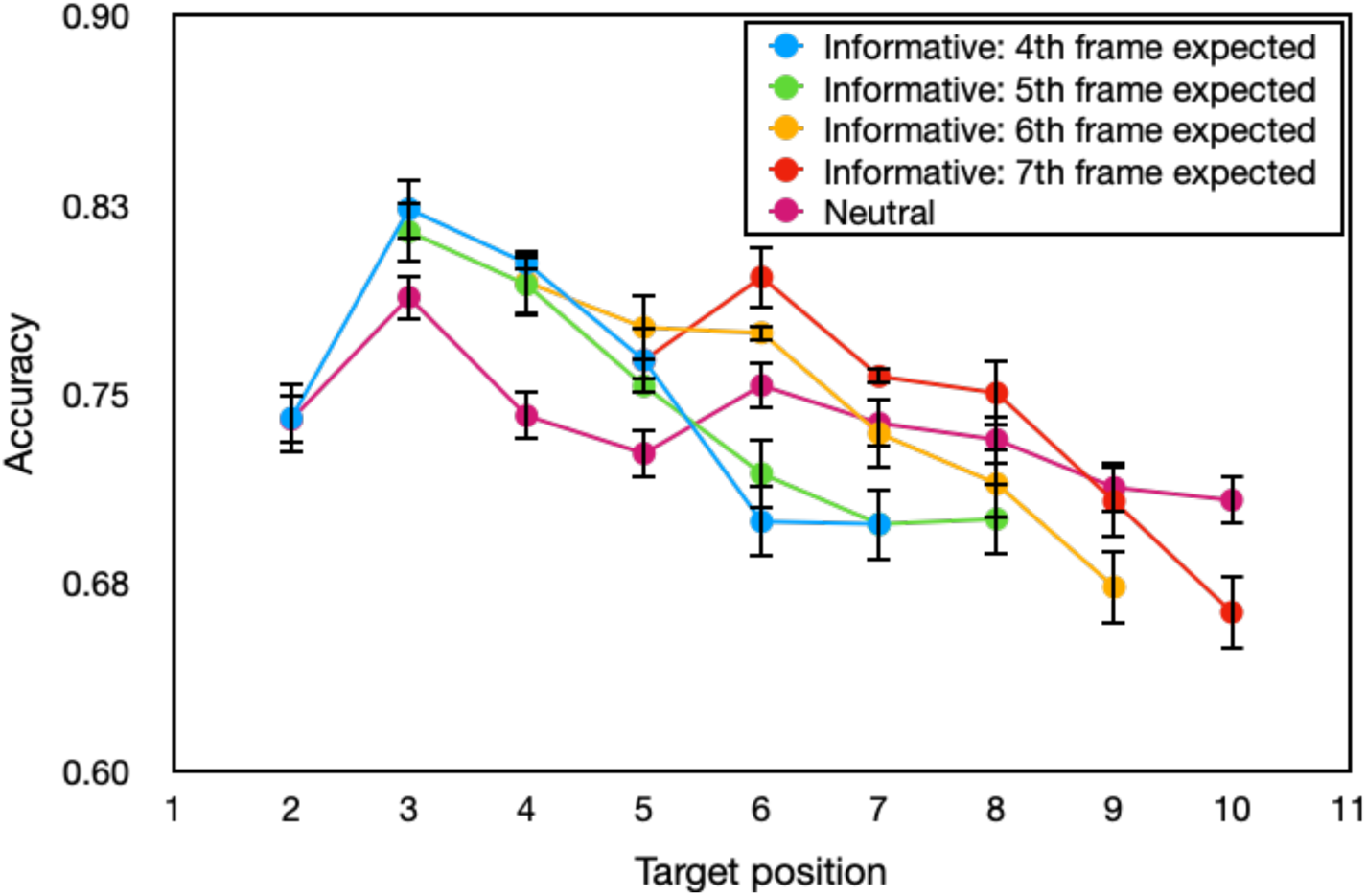
Accuracy in Experiment 4 as a function of informativeness and target position in the sequence of stimuli. Error bars correspond to one SEM across observers.

**Figure 12.**
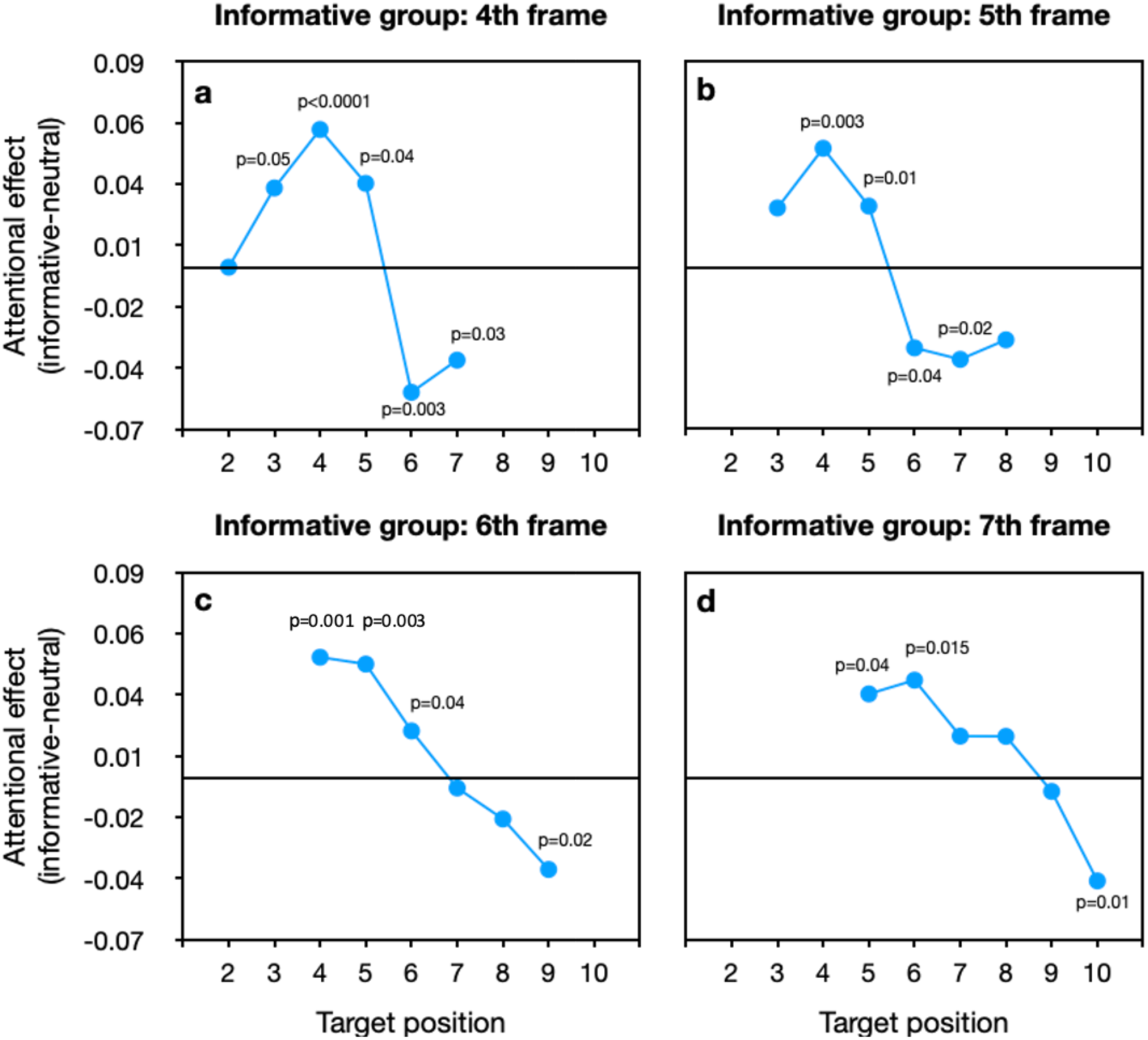
The attentional effect in Experiment 4 as a function of target position within the stream. (a) informative group in which the 4th stimulus position was the expected frame; (b) informative group in which the 5th stimulus position was the expected frame; (c) informative group in which the 6th stimulus position was the expected frame; (d) informative group in which the 7th stimulus position was the expected frame. The p-value for each LSD post-hoc test of the difference between the neutral group and an informative group for a given target position is presented above that target position.

## Discussion

This study examined whether allocating endogenous attention to a particular point in time results in suppression in subsequent time points. Previously, Cutzu and Tsotsos (2003) have demonstrated that attending to a particular location leads to suppression at nearby locations. In their study, 12 items appeared on a circle centered on the fixation point. A precue indicated a location, and the distance of a probed item to the cued location varied. They found that as the distance between the cued location and the probed item increased, the accuracy increased, suggesting that attention is suppressed around the attended location. In a similar task, Yoo et al. (2018) have shown that shifting attention to a particular feature suppresses surrounding features. In their task, the items were oriented bars, and the task was to indicate whether the cued bar and the probed bar were the same or different. They found that the orientation distance between the cued bar and the probed bar affected performance, accuracy was lowest when the orientations were similar, suggesting that attention is suppressed around the attended feature. The current study examined whether attentional suppression also occurs in the time domain. According to the ST model, endogenous attention manipulates the visual processing hierarchy before the target appears so that the visual processing network is ready for the particular moment of target appearance. We designed a paradigm that generates anticipation for the moment the target appears. In each informative block, the target was expected to appear at some point in time, each block different, whereas in the neutral block, the target’s appearance time was unpredictable. Experiments 1-3 revealed effects of block presentation order on performance and suggested a transfer of expectations from informative blocks to the neutral block. Once we eliminated carryover effects in Experiment 4, we found that attention to the expected point in time leads to improved performance at that time and is accompanied by an impaired performance at succeeding time points. This finding supports the general prediction of the ST model regarding the suppression of interfering signals in the hierarchy, that selecting a particular point in time induces suppression at adjacent time points. The suppression of stimulation in the moments after the attended moment (because the past cannot be modulated) reduces the influences of forthcoming distractions, thus enabling more accurate recognition of the stimulus that appeared at the attended time.

This finding is also consistent with spatiotemporal receptive fields’ response properties, characterized by signal enhancement in the early temporal band of cell response and inhibition in the later (Lindeberg, 2021). Studies have demonstrated that spatial and feature-based attention modulate the response properties of the receptive field, such that the center-surround antagonism near the focus of attention is strengthened (e.g., Anton-Erxleben et al., 2009). The current study suggests that temporal attention modifies the receptive field’s properties as well, such that the inhibition of the cell response is strengthened just after the attended time. Recently, Van Ede et al. (2018) investigated the impact of anticipation on both target and distractor representations. Neural activity was recorded while participants observed a Gabor target, after which a distractor could appear. In 50% of the trials, an auditory precue predicted the onset time of the target (i.e., 50% cued trials and 50% uncued trials). At the end of the trial, participants had to reproduce the orientation of the Gabor target. They found a larger distractor interference in uncued trials than in cued trials, so that anticipation improved target representation. Strikingly, looking deeper into the time course of distractor interference on target decoding, they found that the onset of distractor interference was delayed in cued trials. Thus the enhanced target representation might be due to the extended time window that allowed the processing of task-relevant target information without distractor interference. These results are consistent with the ST model and the behavioral performance found in the present study.

The results of Experiments 1-3 suggest that there were carryover effects due to learned probabilities. These results are consistent with Los et al. (2017), who demonstrated that previous timing experiences of target appearance shaped performance, even when a whole week elapsed (Mattiesing et al., 2017). Los et al. (2017) divided the participants into two groups. In the first group, the probability of the target appearing after a short interval from the warning signal (this interval is called foreperiod) was high. While in the second group, the probability of a short foreperiod was low. They found that in the first group, the response time (RT) as a function of foreperiod was flat, whereas, in the second group, RT decreased with increasing foreperiods. Critically, after completing this phase, both groups performed the same block in which all foreperiods varied randomly and with equivalent probable. Strikingly, the RT-foreperiod function for this random block was flatter in the first group than in the second group, demonstrating the influence of previous timing experiences on temporal preparation. Because we compared performance in the informative blocks with performance in the neutral blocks, the transfer effects from the informative to the neutral blocks likely contaminated the results. By using a between-subjects design in Experiment 4, in which each group performed a different condition, we successfully overcame these carryover effects.

Our finding that accuracy improves around the expected temporal position further supports previous demonstrations of improved performance at the attended time point (e.g., Cravo et al., 2011; Niemi & Näätänen, 1981; Yeshurun & Tkacz-Domb, 2021), but the paradigm we used here was different. While in earlier studies, a warning signal indicated the time point of target appearance, and the interval between them was blank, in our study, the target was embedded within a sequence of non-target stimuli. Thus, we replicated previous findings of higher accuracy at the attended time, showing that this effect holds with a different paradigm. Moreover, the results suggest temporal suppression. Interestingly, the results indicate that the estimation of the expected time point was not precise, particularly when the target was expected to appear after the longest duration from the onset of the sequence (i.e., 7th frame expected). This is consistent with the finding that when the interval between the warning signal and the target is fixed throughout the block (i.e., the constant-foreperiod paradigm), performance declines as this interval lengthens (Klemmer, 1956; Niemi, & Näätänen, 1981). In our study, the interval from the beginning of the sequence until the expected time point was not constant during the block, but it was the most probable interval. Still, accuracy in estimating the expected time decreased for the longest interval. It is possible that there is a temporal limitation on how effectively expectation for a target’s appearance can be maintained following a cue (perhaps a short-term memory issue). Specifically, if the interval between the cue and the anticipated time of target appearance becomes too long, the cue’s effectiveness in directing attention to the expected time point may diminish. Consequently, this could result in reduced attentional enhancement and a weaker subsequent suppressive wake. An interesting direction for future research could be to investigate whether generating expectation for a target appearing closer in time to the cue than employed in the present study (e.g., at the 2nd or 3rd frame in the sequence) leads to stronger attentional effects. We did observe stronger suppressive effects for the temporally shorter cues.

Finally, we hypothesized that there would be an orderly movement of the suppression that would follow the expected time point, but the results are noisy. A more robust ordering might be seen if it were possible to test different SOA durations as well as all combinations of ISI and stimulus durations. It is also possible that the time of the visual information to traverse the visual hierarchy is not precisely the same in all participants, and this affects the results in this experiment.

To conclude, this is the first study to examine the dynamic of temporal attention when the target is embedded in a long stream of stimuli, and the observers know in real-time to which target they will have to respond. We found that performance improved at the expected time point and was depressed in the time period following the stimulus that appeared at the expected time. This finding supports the prediction of the ST model that there is a suppression of interfering signals due to attention. In contrast to the many previous examples of this principle where suppression was found in surround regions, here due to the directionality of time, the suppression only follows the expected stimulus. Suppression that occurs of stimuli that follow an expected time point t, means that any interference during the interpretation period is subdued, leading to a clearer perception of the stimulus seen at the expected time point. Such an attention-driven temporal suppression may play a role in the precise timing required for many sports or other dynamic visual behaviors. For example, consider how one might try to return a lob ball in tennis. It seems quite reasonable that your eyes are fixated on the ball and that your attention is as well. Whatever mechanisms ST would have for attending and localizing are locked in. You track the ball in the air against a cluttered background (audience, stadium roof, lights, etc.) in order to time your strike precisely. Not only can you attend spatially and suppress spatial interference, you can also suppress color interference (you know the ball color). You need to precisely measure and track the ball’s speed, acceleration, trajectory, and spin, if not more. These quantities are the domain of the dorsal pathway, and can be quickly computed (Bullier 2001; for the ST motion hierarchy see Tsotsos et al. 2005). Neurons in this pathway necessarily must sample not only space but time as well. They may provide very quick responses but those responses are based on measurements that are evaluated during a time interval (*t_a_, t_b_*); one cannot compute a speed otherwise since *t_b_ - t_a_* is in the denominator of speed computation. One may suppose that each neuron has a time center point *t_c_*, *t_a_ < t_c_* < *t_b_*, and collects information before and after from which a speed, etc. can be computed. The quantization of time across all relevant neurons is assumed to be sufficiently fine-grained and it clearly would be overlapping just like spatial neuron fields. ST’s top down mechanisms, as have been described, suppress irrelevant spatial and featural information. Thus, those do not contribute to the computation within the neuron’s receptive field. But what about the temporal interval? Would modulation in time help with signal-to-noise characteristics as well? Any measurement from early in the time interval would likely have been reported already. The information later in this time interval is newer and of more interest. For the current interval, the early part is in effect noise. Suppression of the older events within the time interval would lead to less interference in computation of the newer time interval events. Note that this also means that *t_b_ - t_c_* is long enough for sufficiently accurate computation. ST’s top-down attentive mechanisms would effect such a suppressive wake together with the other top-down modulations in order to improve the quality and precision of visual motion computations, enabling a precise return of that lob ball.

## Data availability

The data that support the findings of this study are available in the Open Science Framework at https://osf.io/7juxm/?view_only=22210f032a0a46ac9a0ee449ed72275c

## Acknowledgements

This research was funded by Air Force Office of Scientific Research (FA9550-18-1-0054), the Canada Research Chairs Program (950-231659), the Natural Sciences and Engineering Research Council of Canada (RGPIN-2016-05352) grant to J.K.T; and the Israeli Council for Higher Education - Planning and Budgeting Committee grant to S.T.-D.

## References

Anton-Erxleben, K., Stephan, V. M., & Treue, S. (2009). Attention reshapes center-surround receptive field structure in macaque cortical area MT. Cerebral cortex, 19(10), 2466–2478.

Baumeister, A. A., & Joubert, C. E. (1969). Interactive effects on reaction time of preparatory interval length and preparatory interval frequency. Journal of Experimental Psychology, 82(2), 393.

Bartsch, M. V., Loewe, K., Merkel, C., Heinze, H. J., Schoenfeld, M. A., Tsotsos, J. K., & Hopf, J. M. (2017). Attention to color sharpens neural population tuning via feedback processing in the human visual cortex hierarchy. Journal of Neuroscience, 37(43), 10346–10357.

Blom, T., Feuerriegel, D., Johnson, P., Bode, S., & Hogendoorn, H. (2020). Predictions drive neural representations of visual events ahead of incoming sensory information. Proceedings of the National Academy of Sciences, 117(13), 7510–7515.

Boehler, C. N., Tsotsos, J. K., Schoenfeld, M. A., Heinze, H. J., & Hopf, J. M. (2009). The center-surround profile of the focus of attention arises from recurrent processing in visual cortex. Cerebral Cortex, 19(4), 982–991.

Boehler, C. N., Tsotsos, J. K., Schoenfeld, M. A., Heinze, H. J., & Hopf, J. M. (2011). Neural mechanisms of surround attenuation and distractor competition in visual search. Journal of Neuroscience, 31(14), 5213–5224.

Bullier, J. (2001). Integrated model of visual processing. Brain research reviews, 36(2-3), 96–107.

Caputo, G., & Guerra, S. (1998). Attentional selection by distractor suppression. Vision research, 38(5), 669–689.

Carrasco, M. (2011). Visual attention: The past 25 years. Vision research, 51(13), 1484–1525.

Correa, Á., Lupiáñez, J., & Tudela, P. Í. O. (2005). Attentional preparation based on temporal expectancy modulates processing at the perceptual level. Psychonomic bulletin & review, 12(2), 328–334.

Cravo, A. M., Rohenkohl, G., Wyart, V., & Nobre, A. C. (2011). Endogenous modulation of low frequency oscillations by temporal expectations. Journal of neurophysiology, 106(6), 2964–2972.

Cui, X., Stetson, C., Montague, P. R., & Eagleman, D. M. (2009). Ready… go: Amplitude of the FMRI signal encodes expectation of cue arrival time. PLoS biology, 7(8), e1000167.

Cutzu, F., & Tsotsos, J. K. (2003). The selective tuning model of attention: psychophysical evidence for a suppressive annulus around an attended item. Vision research, 43(2), 205–219.

David, S. V., Hayden, B. Y., Mazer, J. A., & Gallant, J. L. (2008). Attention to stimulus features shifts spectral tuning of V4 neurons during natural vision. Neuron, 59(3), 509–521.

DeAngelis, G. C., & Anzai, A. (2004). A modern view of the classical receptive field: Linear and non-linear spatio-temporal processing by V1 neurons. The visual neurosciences, 1, 704–719.

DeAngelis, G. C., Ohzawa, I., & Freeman, R. D. (1995). Receptive-field dynamics in the central visual pathways. Trends in neurosciences, 18(10), 451–458.

Denison, R. N., Carrasco, M., & Heeger, D. J. (2021). A dynamic normalization model of temporal attention. Nature human behaviour, 5(12), 1674–1685.

Denison, R. N., Heeger, D. J., & Carrasco, M. (2017). Attention flexibly trades off across points in time. Psychonomic bulletin & review, 24(4), 1142–1151.

Faul, F., Erdfelder, E., Lang, A. G., & Buchner, A. (2007). G* Power 3: A flexible statistical power analysis program for the social, behavioral, and biomedical sciences. Behavior research methods, 39(2), 175–191.

Fischer, J., & Whitney, D. (2009). Attention narrows position tuning of population responses in V1. Current biology, 19(16), 1356–1361.

Ghose, G. M., & Maunsell, J. H. (2002). Attentional modulation in visual cortex depends on task timing. Nature, 419(6907), 616–620.

Griffin, I. C., Miniussi, C., & Nobre, A. C. (2001). Orienting attention in time. Frontiers in Bioscience, 6(12), D660–D671.

Haenny, P. E., & Schiller, P. H. (1988). State dependent activity in monkey visual cortex. Experimental Brain Research, 69(2), 225–244.

Klemmer, E. T. (1956). Time uncertainty in simple reaction time. Journal of experimental psychology, 51(3), 179.

Li, Q., Joo, S. J., Yeatman, J. D., & Reinecke, K. (2020). Controlling for participants’ viewing distance in large-scale, psychophysical online experiments using a virtual chinrest. Scientific reports, 10(1), 1–11.

Lindeberg, T. (2021). Normative theory of visual receptive fields. Heliyon, 7(1), e05897.

Los, S. A., Kruijne, W., & Meeter, M. (2017). Hazard versus history: Temporal preparation is driven by past experience. Journal of Experimental Psychology: Human Perception and Performance, 43(1), 78.

Martinez-Trujillo, J. C., & Treue, S. (2004). Feature-based attention increases the selectivity of population responses in primate visual cortex. Current biology, 14(9), 744–751.

Mattiesing, R. M., Kruijne, W., Meeter, M., & Los, S. A. (2017). Timing a week later: The role of long-term memory in temporal preparation. Psychonomic bulletin & review, 24(6), 1900–1905.

Mehta, A. D., Ulbert, I., & Schroeder, C. E. (2000). Intermodal selective attention in monkeys. I: distribution and timing of effects across visual areas. Cerebral cortex, 10(4), 343–358.

Moon, J., Choe, S., Lee, S., & Kwon, O. S. (2019). Temporal dynamics of visual attention allocation. Scientific reports, 9(1), 1–11.

Müller, N. G., Mollenhauer, M., Rösler, A., & Kleinschmidt, A. (2005). The attentional field has a Mexican hat distribution. Vision research, 45(9), 1129–1137.

Niemi, P., & Näätänen, R. (1981). Foreperiod and simple reaction time. Psychological bulletin, 89(1), 133.

O’Connor, D. H., Fukui, M. M., Pinsk, M. A., & Kastner, S. (2002). Attention modulates responses in the human lateral geniculate nucleus. Nature neuroscience, 5(11), 1203–1209.

Peirce, J., Gray, J. R., Simpson, S., MacAskill, M., Höchenberger, R., Sogo, H., Kastman, E., & Lindeløv, J. K. (2019). PsychoPy2: Experiments in behavior made easy. Behavior research methods, 51(1), 195–203.

Reynolds, J. H., & Heeger, D. J. (2009). The normalization model of attention. Neuron, 61(2), 168–185.

Roberts, M., Delicato, L. S., Herrero, J., Gieselmann, M. A., & Thiele, A. (2007). Attention alters spatial integration in macaque V1 in an eccentricity-dependent manner. Nature neuroscience, 10(11), 1483–1491.

Rohenkohl, G., Cravo, A. M., Wyart, V., & Nobre, A. C. (2012). Temporal expectation improves the quality of sensory information. Journal of Neuroscience, 32(24), 8424–8428.

Posner, M. I. (1980). Orienting of attention. Quarterly journal of experimental psychology, 32(1), 3–25.

Thorpe, S., Fize, D., & Marlot, C. (1996). Speed of processing in the human visual system. nature, 381(6582), 520–522.

Tombu, M., & Tsotsos, J. K. (2008). Attending to orientation results in an inhibitory surround in orientation space. Perception & Psychophysics, 70(1), 30–35.

Tsotsos, J. K. (1988). A ‘complexity level’ analysis of immediate vision. International Journal of Computer Vision, 1(4), 303–320.

Tsotsos, J. K. (1990). Analyzing vision at the complexity level. Behavioral and brain sciences, 13(3), 423–445.

Tsotsos, J. K., Liu, Y., Martinez-Trujillo, J. C., Pomplun, M., Simine, E., & Zhou, K. (2005). Attending to visual motion. Computer Vision and Image Understanding, 100(1-2), 3–40.

Tsotsos, J. K., Rodriguez-Sanchez, A. J., Rothenstein, A. L., & Simine, E. (2008). Different binding strategies for the different stages of visual recognition. Brain Research, 1225, 119–132.

Tsotsos, J. K. (2011). A computational perspective on visual attention. MIT Press.

van Ede, F., Chekroud, S. R., Stokes, M. G., & Nobre, A. C. (2018). Decoding the influence of anticipatory states on visual perception in the presence of temporal distractors. Nature communications, 9(1), 1–12.

Womelsdorf, T., Anton-Erxleben, K., Pieper, F., & Treue, S. (2006). Dynamic shifts of visual receptive fields in cortical area MT by spatial attention. Nature neuroscience, 9(9), 1156–1160.

Yeshurun, Y., & Tkacz-Domb, S. (2021). The time-course of endogenous temporal attention–Super fast voluntary allocation of attention. Cognition, 206, 104506.

Yoo, S. A., Tsotsos, J. K., & Fallah, M. (2018). The attentional suppressive surround: Eccentricity, location-based and feature-based effects and interactions. Frontiers in Neuroscience, 710.

Yoo, S. A., Martinez-Trujillo, J. C., Treue, S., Tsotsos, J. K., & Fallah, M. (2021). Feature-based attention induces surround suppression during the perception of visual motion. bioRxiv, 2021-02.

